# rt-me-fMRI: A task and resting state dataset for real-time, multi-echo fMRI methods development and validation

**DOI:** 10.1101/2020.12.07.414490

**Authors:** Stephan Heunis, Marcel Breeuwer, César Caballero-Gaudes, Lydia Hellrung, Willem Huijbers, Jacobus FA Jansen, Rolf Lamerichs, Svitlana Zinger, Albert P Aldenkamp

## Abstract

A multi-echo fMRI dataset (N=28 healthy participants) with four task-based and two resting state runs was collected, curated and made available to the community. Its main purpose is to advance the development of methods for real-time multi-echo functional magnetic resonance imaging (rt-me-fMRI) analysis with applications in neurofeedback, real-time quality control, and adaptive paradigms, although the variety of experimental task paradigms supports a multitude of use cases. Tasks include finger tapping, emotional face and shape matching, imagined finger tapping and imagined emotion processing. This work provides a detailed description of the full dataset; methods to collect, prepare, standardize and preprocess it; quality control measures; and data validation measures. A web-based application is provided as a supplementary tool with which to interactively explore, visualize and understand the data and its derivative measures: https://rt-me-fmri.herokuapp.com/. The dataset itself can be accessed via a data use agreement on DataverseNL at https://dataverse.nl/dataverse/rt-me-fmri. Supporting information and code for reproducibility can be accessed at https://github.com/jsheunis/rt-me-fMRI.

## 1. Background and summary

Real-time functional magnetic resonance imaging (fMRI) is a brain imaging method where functional brain signals are acquired, processed, and used during an ongoing scanning session. Applications include real-time data quality control (Dosenbach et al., 2017), adaptive experimental paradigms (Hellrung et al., 2015), and neurofeedback (Sitaram et al., 2017). Neurofeedback is a cognitive training method where the real-time feedback signal is presented back to the participant to allow self-regulation of their blood oxygen level-dependent (BOLD) signal, prompting researchers to investigate it as an intervention for patients with neurological or psychiatric conditions. Work by Ros et al. (2020) and Haugg et al. (2020) show an absence of standardisation in experimental design and outcome reporting restricts the synthesis of evidence to determine the efficacy of fMRI neurofeedback. Further, it remains a major challenge to delineate the sources of variance in the brain and in neurofeedback signals and their eventual effects on neurofeedback training outcomes. Similar challenges exist for separating BOLD and non-BOLD variations and their influences on data quality, and subsequently on all real-time fMRI applications.

In recent work (Heunis et al., 2020a) we investigated the available acquisition and processing methods for improving real-time fMRI signal quality, and identified an absence of methodological denoising studies and a need for community-driven quality control standards. Here, we aim to advance this process by curating a multi-echo fMRI dataset (***rt-me-fMRI***). It builds on known benefits of multi-echo fMRI for increasing BOLD sensitivity both in resting state and task fMRI (Olafsson et al., 2015; Gonzalez-Castillo et al., 2016; Kundu et al., 2017; Dipasquale et al., 2017; Moia et al., 2020). Potential benefits of multi-echo fMRI in the real-time context have been reported before (Posse et al., 2000; Posse et al., 2003; Weiskopf et al., 2005; Marxen et al., 2016), but real-time multi-echo processing methods remain underexplored. By releasing the ***rt-me-fMRI*** dataset, we aim to facilitate a community effort to advance the development of methods and standards in this domain.

The ***rt-me-fMRI*** dataset includes multi-echo resting state and task-based fMRI data from 28 healthy participants. Fig.1 provides an overview, including the task types: finger tapping, emotion processing, imagined finger tapping, and imagined emotion. Several factors influenced the experimental and acquisition protocols:

***Multi-echo fMRI***: To facilitate the development of real-time multi-echo methods, all functional acquisitions have multiple echoes. The first resting state run allows calculation of quantitative multi-echo parameters such as baseline *T_2_*\* or *S_0_* maps, which can in turn be used for echo combination during subsequent runs.
***Task and resting state:*** The motor cortex, amygdala, and visual system were selected as representative regions based on frequency of studies in fMRI and neurofeedback literature (Thibault et al., 2018), and tasks were selected to elicit appropriate BOLD responses. The *fingerTappingImagined* and *emotionProcessing* tasks respectively allow investigations into mental imagery and visual shape/face processing. Since these structures are located at distinct anatomical regions that experience different levels of noise (e.g. the amygdala suffers from more severe image dropout and physiological noise; Boubela et al., 2015), this allows investigation of spatially distinct effects of real-time denoising. Resting state scans allow comparison of the effects of processing steps in the absence and presence of a task.
***Template data:*** In real-time fMRI applications, anatomical and functional scans are typically acquired before the main session to generate registration, segmentation, and localisation templates. This assists real-time realignment and extraction of region-based signals, and minimises per-volume processing time.
***No neurofeedback:*** To keep the setup applicable to a range of real-time scenarios without introducing additional confounds, no neurofeedback was provided. Instead, to approximate similar mental states, the second functional set of scans were structured as imagined versions of the first functional set. This is a common approach in neurofeedback training: amygdala neurofeedback participants have been asked to think about an emotional event in their past (e.g. Young et al., 2014; Misaki et al., 2018), while motor cortex neurofeedback participants have been asked to think about performing physical exercises (e.g. Subramanian et al., 2011).
***Physiology data:*** To facilitate the development and exploration of real-time physiological denoising methods and their relation to multi-echo-derived data, cardiac and respiratory signals were acquired.

**Figure 1:**
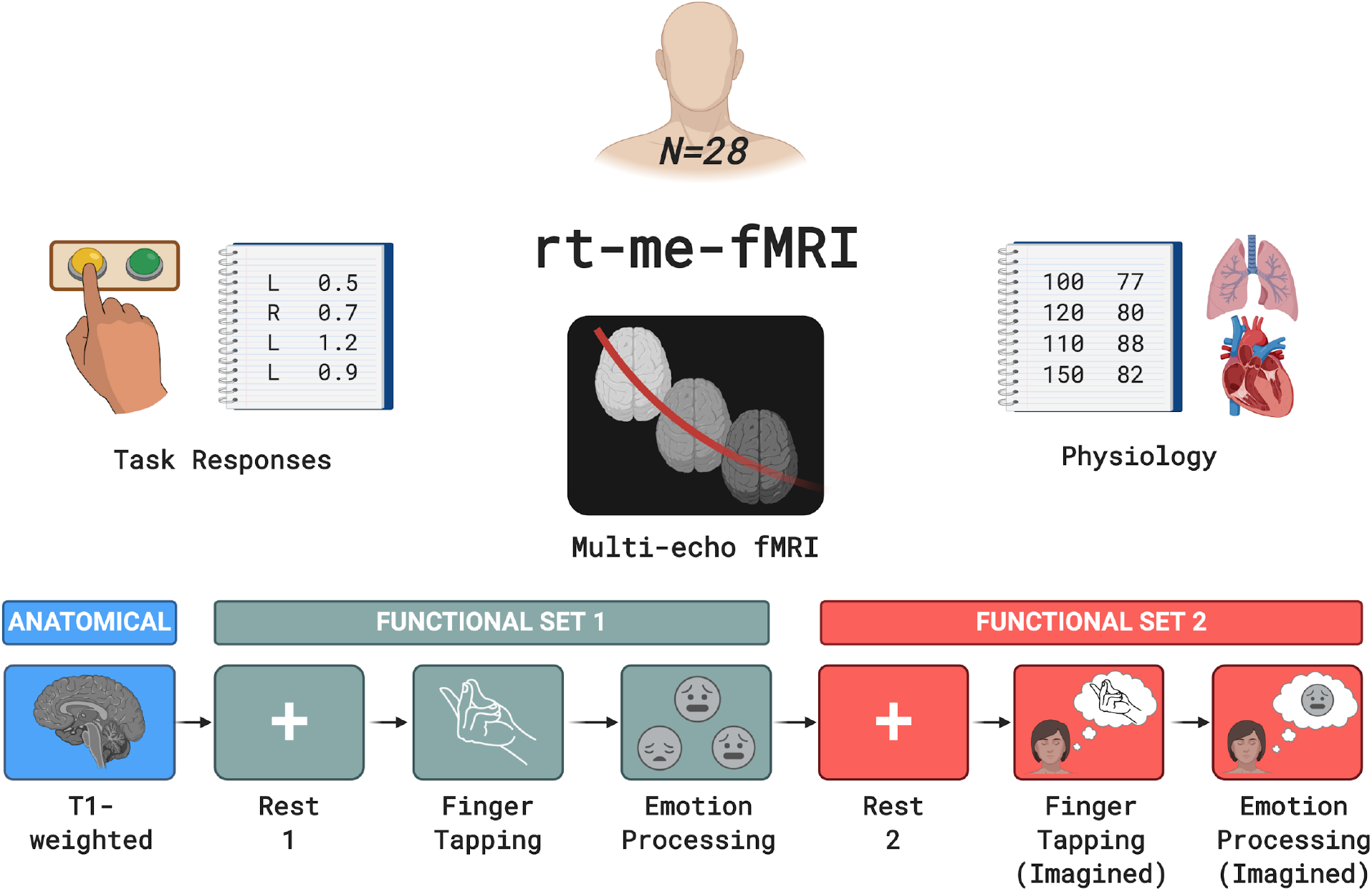
A depiction of the rt-me-fMRI dataset collected for 28 healthy participants. Acquired data include anatomical MRI, resting state and task-based multi-echo fMRI, task responses and physiology data. The bottom row indicates the order and type of acquired MRI scans. Colour-coding separates the anatomical scan from functional set 1 and from functional set 2. Functional set 1 includes resting state, fingerTapping and emotionProcessing acquisitions, while functional set 2 includes resting state, fingerTappingImagined, and emotionProcessingImagined acquisitions.

The ***rt-me-fMRI*** is available in BIDS format via the DataverseNL repository: https://dataverse.nl/dataverse/rt-me-fmri. A browser-based environment allows interactive exploration of the data quality and derivatives (https://rt-me-fmri.herokuapp.com/).

## 2. Methods

### 2.1. Ethics and data privacy

The data described here was collected as part of a study for which ethics approval was granted by two ethics review boards. To confirm that the study protocol is in accordance with the Dutch national law on medical-scientific research conducted on human participants (see WMO: https://wetten.overheid.nl/BWBR0009408/2020-01-01), the medical ethical review board at the Máxima Medisch Centrum (Veldhoven, NL) granted ethics approval. Secondly, the local ethics review board at Kempenhaeghe Epilepsy Center (Heeze, NL; where the data was collected) approved the study protocol.

All participants provided informed and written consent to participate in the study and for their maximally de-identified data (also referred to as limited data) to be shared publicly under specific conditions (see GDPR considerations below). Participants were provided with an electronic version of a “Participant Information Letter” which contained, in addition to standard information about the study protocol, clear information about their personal data privacy and the risks and benefits involved in sharing maximally de-identified versions of their data. They were asked to read it thoroughly and to discuss it with friends and family if they wished to do so. They were granted an opportunity to discuss any questions or concerns about their voluntary participation in the study with the lead researcher, both via email and in person. If they decided to continue with participation, participants signed the consent form and were provided with an electronic copy.

The dataset was collected, processed and shared in accordance with the European Union’s General Data Protection Regulation (GDPR) as approved by Data Protection Officers (DPOs) at Kempenhaeghe Epilepsy Center (Heeze, NL) and the Eindhoven University of Technology. Of particular note is the procedure that was followed to enable sharing of the dataset under specific conditions that allow personal data privacy to be prioritised while adhering to FAIR data standards (“findable, accessible, interoperable, reusable”; see Wilkinson et al., 2016), with this being the first documented implementation. It followed from the collaborative effort of the Open Brain Consent Working Group (Pernet et al., 2020), a group of researchers, data experts, and legal practitioners that aim to provide globally standardised templates for informed consent and data privacy statements that allow for brain research data to be shared while prioritising personal data privacy. Steps to accomplish this include following best practices to de-identify brain images (e.g. removing personally identifiable information from image filenames and metadata and removing facial features from T1-weighted images), converting the data to BIDS format, employing a Data Use Agreement, and keeping participants fully informed about each of these steps and the associated risks and benefits. The Data Use Agreement can be accessed in this manuscript’s GitHub repository: https://github.com/jsheunis/rt-me-fMRI.

### 2.2. Participants

The ***rt-me-fMRI*** dataset consists of MRI and physiology data from 28 healthy, right-handed (self-report) adults recruited from the local student population: 20 male, 8 female; age = 24.9 ± 4.7 (mean ± standard deviation). During recruiting, possible participants were excluded if they reported prior or current (at the time of the study) indications of neurological or psychiatric conditions, or any other standard contraindications for MRI scanning. 31 participants were initially recruited for the dataset, but three were excluded because of technical and administrative challenges. All anatomical scans were inspected by a trained radiologist and no incidental findings were reported.

### 2.3. Experimental protocol

#### 2.3.1. Preparation and instructions

A single experimenter interacted with all participants. Data for each participant was collected during a single scanning session of approximately 1 hour, preceded by a 30 min onboarding procedure and followed by a 15 min offboarding procedure. Onboarding included a tour of the scanner and related equipment, detailed instructions for the participant to follow during each scan, and time for additional questions.

To minimise participant motion during scans so as to improve spatial and temporal image quality, participants were asked to remain as still as possible inside the scanner. Additionally, a length of tape was fixed across the participants’ foreheads to the stationary part of the head coil. This provided tactile feedback which has been demonstrated as a simple and effective way to reduce head motion during fMRI scanning (Krause et al., 2019).

Lights in the scanner room were dimmed during the experiment. Participants viewed instructions projected on a screen at the back of the scanner bore via a head coil-mounted mirror. For resting state functional scans, participants were instructed to keep their eyes open and fixate on the cross on the screen.

#### 2.3.2. Experimental design

All functional scans have 210 volumes and exactly the same sequence parameters. All task scans follow a block design with 10 volumes (i.e. 20 s) per block, and with blocks alternating between control and task conditions. All task designs start and end with a control condition. These block design aspects are depicted in Fig. 2 below for all task runs. Take note that the depictions do not necessarily agree with the exact stimuli as seen by the participants, as the depictions below are purely illustrative.

**Figure 2:**
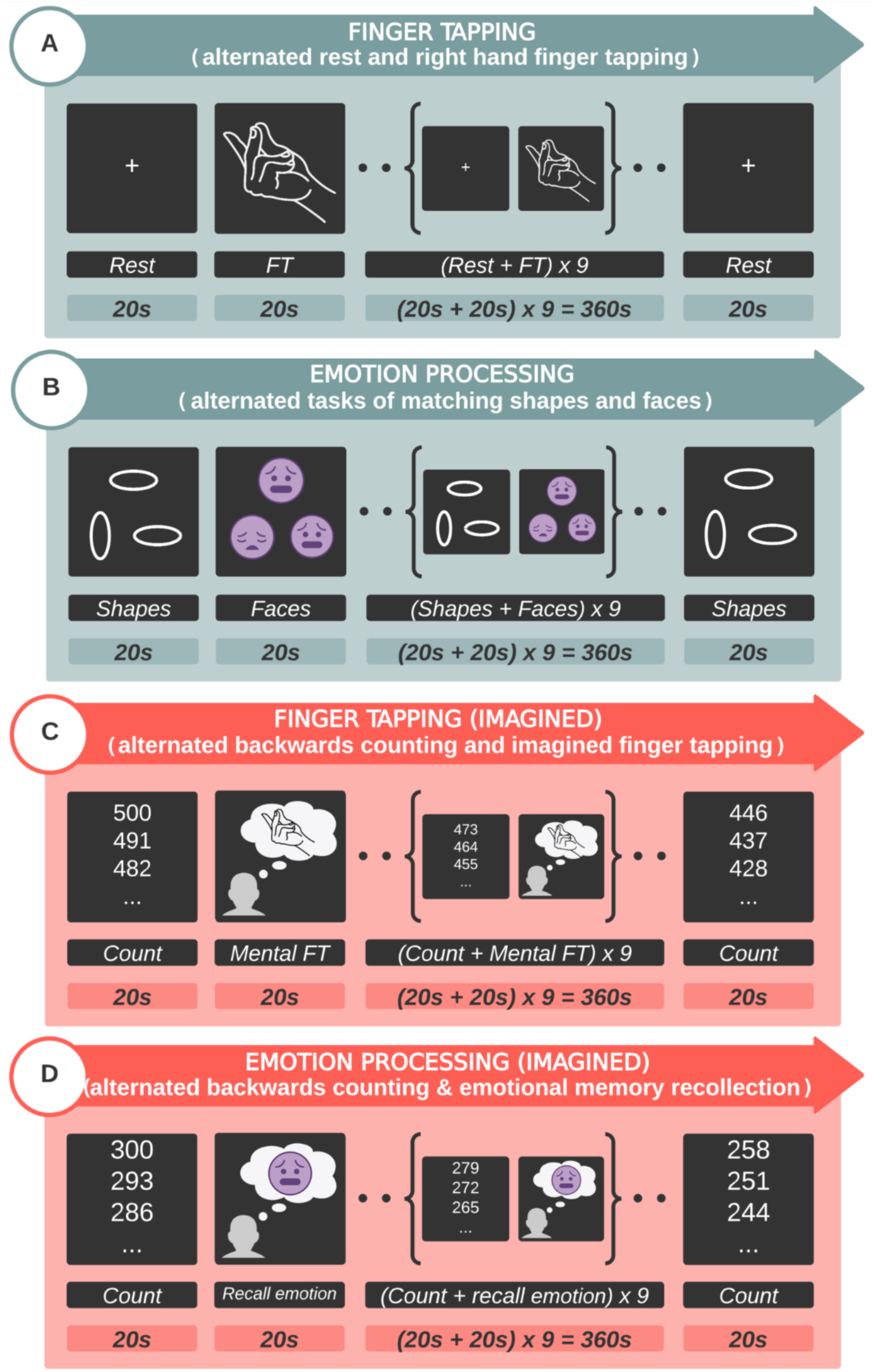
Depictions of the experimental designs for all tasks. Subfigures include: (A) fingerTapping - right hand finger tapping, (B) emotionProcessing - matching shapes and faces, (C) fingerTappingImagined - imagined finger tapping, and (D) emotionProcessingImagined - emotional memory recollection. All designs follow a block paradigm with 10 volumes (i.e. 20 s) per block, and with blocks alternating between control and task conditions. All task designs start and end with a control condition. Color code: functional set 1 = Green; functional set 2 = Red. FT = finger tapping.

For the *fingerTapping* task, participants were instructed to execute finger tapping with their right hand by steadily tapping the tip of the thumb to the tip of each other finger in succession, reversing the tapping order until the end of the task block is reached. For the *fingerTappingImagined* task, participants were instructed to imagine doing exactly the same as in the actual finger tapping task, but without actually moving their right fingers. For the control condition during the *fingerTappingImagined* task, participants were asked to count backwards in multitudes of 7.

The *emotionProcessing* task was an adapted “Hariri” task from the emotion processing task used in the Human Connectome Project (Van Essen et al., 2013; Hariri et al., 2002; Manuck et al., 2007). Materials were implemented to suit the paradigm for this ***rt-me-fMRI*** dataset. During each 20 s task block, participants were presented with a task cue (3 s duration), followed by a trial with three pictures of faces where the participant had to select one of the bottom figures (left or right) that resembled the top one, by pressing a left or right button (2 s duration). The inter-trial interval was 1 s duration (see Fig. 3). Each 20 s block had 6 trials. The same design timing was used for the control condition blocks, i.e. matching shapes, as for the trial condition blocks depicted in Fig.

**Figure 3:**
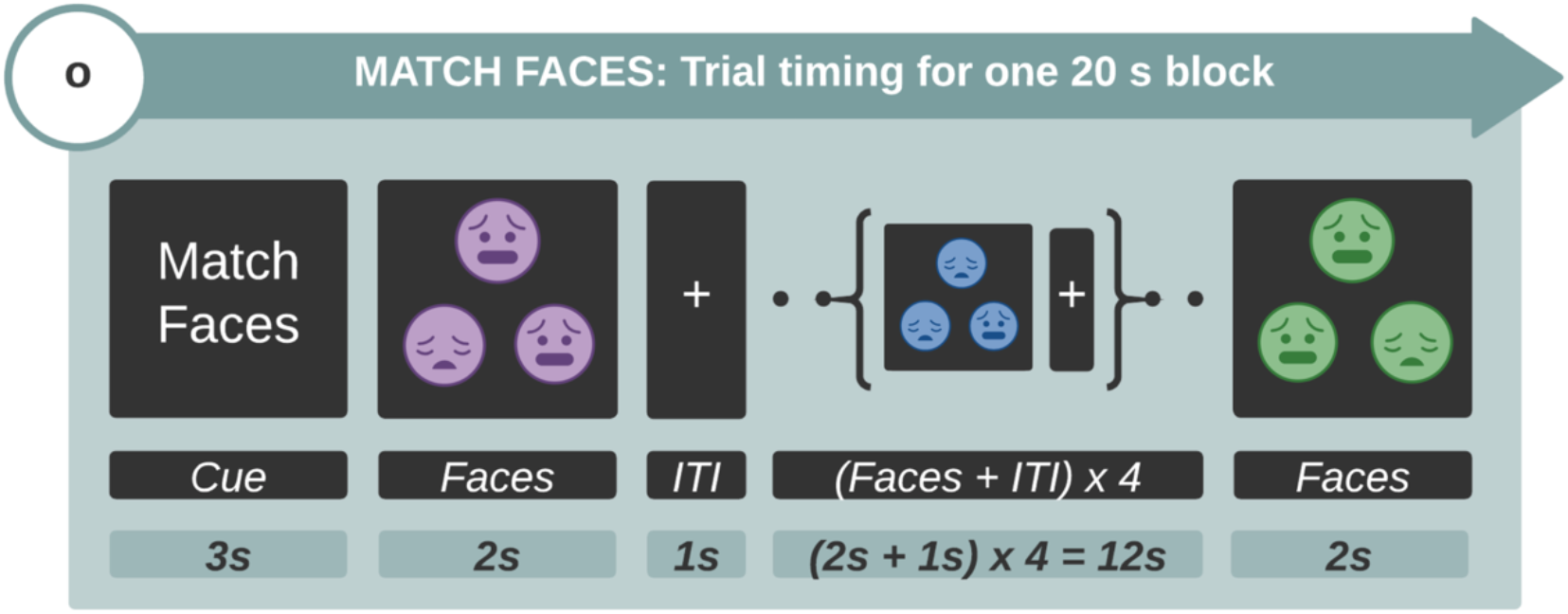
The task timing for the emotionProcessing task. Times are provided for the cue, trials and inter-trial interval during a single block (20 s) of the face matching condition. The same design timing was used for the control condition blocks.

Participants used an MRI-compatible button box with their right hand to complete the task. Participants were asked to press the left button with their right index finger if selecting the bottom left image (shape or face) on the screen, and to conversely press the right button with their right middle finger if selecting the bottom right image.

For the *emotionProcessingImagined* task, participants were instructed prior to the scanning session to identify an emotional event in their past that involved a person or people, and to think about this event and also try to mentally experience the identified emotion during the task blocks. For the control condition during this mental emotion task, participants were asked to count backwards in multitudes of 9.

Participants were interviewed after the scanning session about their experiences during the MRI acquisition and the tasks. None reported detrimental issues with regards to their ability to focus on the task or with task-switching.

Tasks and instructions were programmed and presented to the participants using E-Prime Studio version 2.0.10.248. The programmed E-prime files used for each task (“.es2” format), as well as all presented images for trials, conditions, cues and instructions (.jpg format), can be accessed in the supplementary code repository: https://github.com/jsheunis/rt-me-fMRI. The exact timing information for the presented material (for all functional runs) and the button presses (for *emotionProcessing*), as well as the actual button press responses, were exported from E-prime (in .dat and .txt format) at the end of each session.^1^

### 2.4. MRI acquisition parameters

MRI data was acquired on a 3 Tesla Philips Achieva scanner (software version 5.1.7) and using a Philips 32-channel head coil.

#### 2.4.1. Anatomical MRI

A single T1-weighted anatomical image was acquired using a 3D gradient echo sequence (T1 TFE) with scanning parameters: TR = 8.2 ms; TE = 3.75 ms; flip angle = 8°; field of view = 240×240×180 mm^3^; resolution = 1×1×1 mm^3^; total scan time = 6:02 min.

#### 2.4.2. Functional MRI

All six functional MRI scans were acquired using a multi-echo, echo-planar imaging sequence with scanning parameters: TR = 2000 ms; TE = 14, 28, 42 ms (3 echoes); number of volumes = 210 (excluding 5 dummy volumes discarded by the scanner); total scan time = 7:00 min (excluding 5 dummy volumes); flip angle = 90°; field of view = 224×224×119 mm^3^; resolution = 3.5×3.5×3.5 mm^3^; in-plane matrix size = 64×64; number of slices = 34; slice thickness = 3.5 mm; interslice gap = 0 mm; slice orientation = oblique; slice order/direction = sequential/ascending; phase-encoding direction = A/P; SENSE acceleration factor = 2.5; parts of the cerebellum and brainstem were excluded for some participants to ensure full motor cortex and amygdala coverage.

The echo times, spatial resolution, and SENSE factor were tuned with the aim of improving spatial resolution and coverage while limiting the TR at maximum 2000 ms, including a maximum number of echoes, and keeping the SENSE factor low to prevent SENSE artefacts.

### 2.5. Physiology data acquisition parameters

Breathing fluctuations were recorded with the use of a pressure-based breathing belt strapped around the participant’s upper abdomen. Heart rate was recorded using a pulse oximeter fixed to the participant’s left index finger. Both of these recording devices were wired directly to the scanner, sampled at 500 Hz, synchronized internally to the start/stop pulses of each functional scan, and data were written to Philips’s standard “scanphyslog” log file type.

### 2.6. Standardization: Brain Imaging Data Structure

To adhere to FAIR data principles, the full dataset was curated into the standardized and community-maintained Brain Imaging Data Structure (BIDS; Gorgolewski et al., 2016). This involved the use of several software packages and custom scripts to assist in file format conversion and data structuring, as detailed below. A Jupyter notebook containing Python code and descriptions for each of the steps below can be accessed at the project’s code repository: https://github.com/jsheunis/rt-me-fMRI.

#### 2.7.1. MRI data

Anatomical and functional MRI data were converted from the Philips PAR/REC format to BIDS using the Python package *bidsify* (v0.3; https://github.com/NILAB-UvA/bidsify) This package has *dcm2niix* (v1.0.20190410; https://github.com/rordenlab/dcm2niix/releases/tag/v1.0.20190410) as a dependency to convert the PAR/REC files to NIfTI. It also structures the data into the directory system specified by the BIDS standard.

Anatomical files were additionally de-identified using *pydeface* (v2.0.0; https://github.com/poldracklab/pydeface/releases/tag/2.0.0; Gulban et al., 2019), which removes facial features from the T1w NIfTI image. Further anonymization steps included removing time and date stamps and any identifiable information related to the acquisition location or system from the files output from *bidsify*.

Since PAR/REC files do not contain slice timing information, the converted NIfTI files did not contain it either. Slice timing information was calculated using available parameters and added with a script to the BIDS-specific JSON sidecar files.

#### 2.7.2. Physiology data

Heart rate and breathing traces were converted from the Philips “scanphyslog” format to BIDS format using the Python package *scanphyslog2bids* (v0.1; https://github.com/lukassnoek/scanphyslog2bids).

#### 2.7.3. Task presentation and response data

Presentation timing, button presses and button press response timing information were all converted to the BIDS format using a combination of custom Python scripts and the *convert-eprime* package (v0.0.1; https://github.com/tsalo/convert-eprime/releases/tag/0.0.1; Salo, 2020).

### 2.8. Preprocessing

Raw data was preprocessed using the open source MATLAB-based and Octave-compatible *fMRwhy* toolbox (see “Code Availability” for details). The basic anatomical and functional preprocessing pipeline applied to all data is depicted in Fig. 4 below.

**Figure 4:**
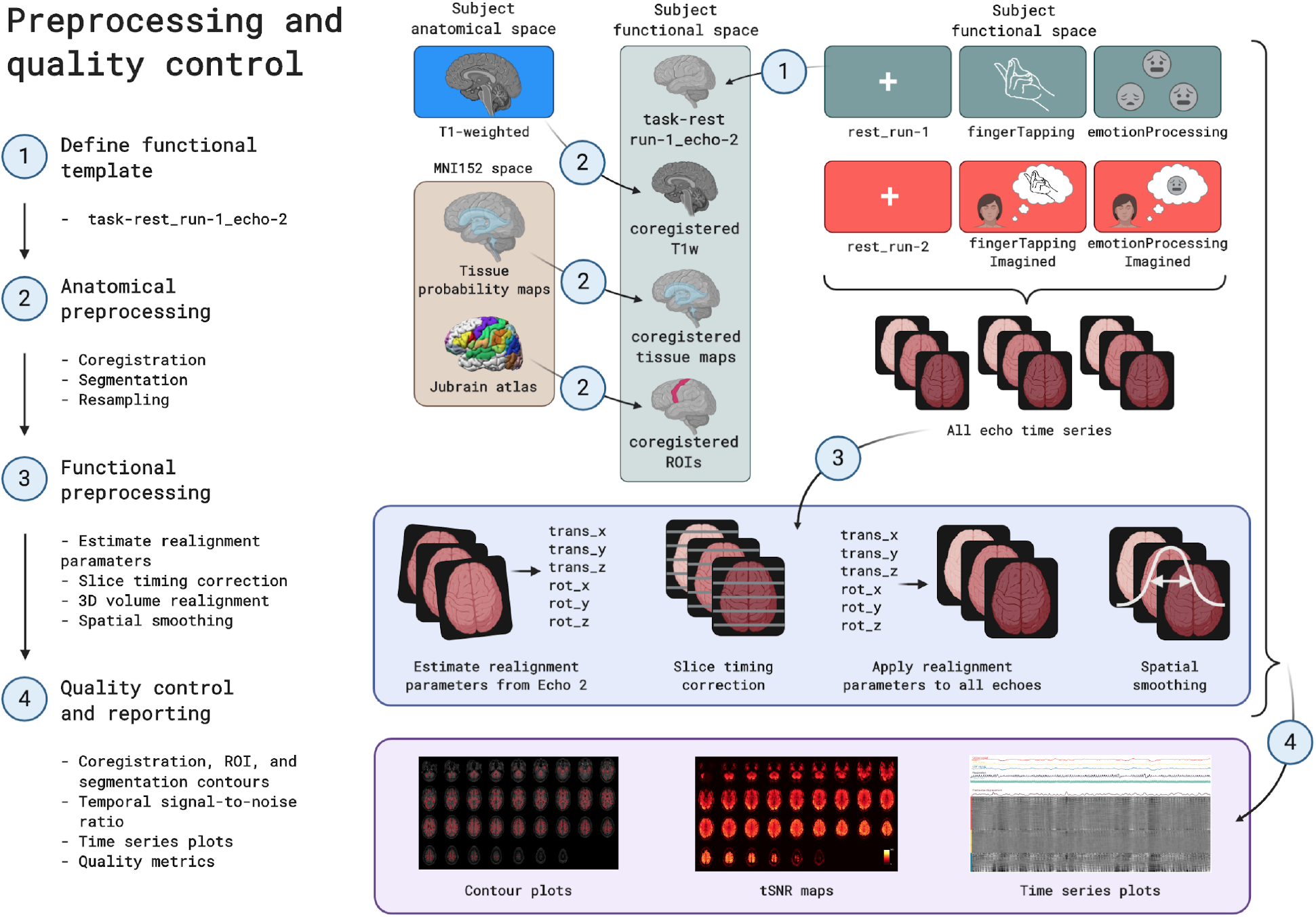
A diagram depicting the preprocessing steps conducted on the rt-me-fMRI dataset in chronological order. Steps include: (1) defining a functional template image from the first resting state run; (2) mapping the anatomical image and atlas-based regions of interest to the functional template space; (3) estimating realignment parameters from the template echo time series, running slice timing correction, applying realignment parameters to all echo time series, and applying spatial smoothing, and (4) generating quality control metrics and visualizations for anatomical and functional data.

As a first step, the T1-weighted anatomical image was coregistered to the template functional image (*task-rest_run-1_echo-2*, volume 1) using SPM12’s *coregister/estimate* functionality, which maximizes normalised mutual information to generate a 12 degree-of-freedom transformation matrix. Before resampling to the functional resolution, this coregistered T1-weighted image was segmented using tissue probability maps and SPM12’s unified segmentation algorithm (Ashburner and Friston, 2005). This yielded subject-specific probability maps for gray matter, white matter, CSF, soft tissue, bone and air in the subject functional space. All of these probability maps were then resampled (using *coregister/write*) to the subject functional resolution. Masks were generated for gray matter, white matter, CSF, and the whole brain (a combination - logical OR after thresholding - of the previous three masks). These were overlaid on the coregistered and resampled T1w image below, to allow visual inspection of segmentation and registration quality.

Anatomical regions of interest were then taken from the cytoarchitecture-based atlases in the SPM Anatomy Toolbox (Eickhoff et al., 2005). For the motor cortex, regions *4a* and *4p* were used. For the amygdala, regions *LB, IF, SF, MF, VTM*, and *CM* were used. For the fusiform gyrus, regions *FG1*, *FG2*, *FG3*, and *FG4* were used. Regions of interest were transformed from MNI152 space to the subject functional space using SPM12 *normalise/write*, as well as the inverse transformation field that was saved as part of the segmentation procedure mentioned above. The regions of interest for this study include the left motor cortex (for the motor processing tasks), the bilateral amygdala (for the emotion processing tasks) and the fusiform gyrus (for the *emotionProcessing* task). These ROIs are overlaid on the coregistered and resampled T1-weighted image, to allow visual inspection of normalisation quality.

Functional data were preprocessed, starting with estimating realignment parameters for each functional time series using SPM12’s *realign/estimate,* which performs a 6 degree-of-freedom rigid body transformation that minimizes the sum of squared differences between each volume and the template volume. Realignment parameters were estimated for the second-echo time series of each run. Then, slice timing correction was done with SPM12, which corrects for differences in image acquisition time between slices. Each echo time series of all functional runs were slice time corrected. 3D volume realignment followed, which applied spatial transformation matrices derived from the previously estimated realignment parameters to all echo time series of all functional runs. Both raw time series and slice time corrected time series were realigned. Lastly, all echo time series of all functional runs were spatially smoothed using a Gaussian kernel filter with FWHM = 7 mm (i.e. double the voxel size). Smoothing was performed on raw, slice time corrected and realigned time series data.

Next, several signal time series were calculated or extracted for use as possible GLM regressors in functional task analysis, or for quality control. From the realignment parameters (3 translation and 3 rotation parameters per volume), a Volterra expansion yielded derivatives, squares and squares of derivatives (Friston et al., 1996). Framewise displacement (FD, Power et al., 2012) was also calculated from the realignment parameters, and volumes were marked as outliers based on different thresholds of, respectively, 0.2 mm and 0.5 mm. RETROICOR regressors (Glover et al., 2000) were generated from the cardiac and respiratory signals using the *TAPAS PhysIO* toolbox, which yielded 6 cardiac regressors, 8 respiratory regressors, 4 interaction regressors, and additionally a cardiac rate regressor (CR; the cardiac rate time series convolved with the cardiac response function; Chang et al., 2009) and a respiratory volume per time regressor (RVT; respiratory volume per time convolved with the respiratory response function; Birn et al. 2006; Birn et al., 2008). From the slice time corrected and realigned time series data (of all functional runs), signals were extracted per voxel and spatially averaged within the previously generated tissue masks to yield tissue compartment signals for gray matter, white matter, cerebrospinal fluid (CSF) and the whole brain.

The last set of preprocessing steps included calculation of image quality metrics and visualizations, using the BIDS-compatible *fmrwhy_bids_workflowQC* pipeline from the *fMRwhy* toolbox. Operations on functional time series data were all done on detrended (linear and quadratic trends) realigned data, except where otherwise specified. Temporal signal-to-noise ratio (tSNR) maps were calculated for all runs by dividing the voxel-wise time series mean by the voxel-wise standard deviation of the time series. Tissue compartment averages were then extracted from these tSNR maps. Percentage difference maps (from the time series mean) were calculated per volume for use in carpet plots (or gray plots).

## 3. Data Records

The ***rt-me-fMRI*** dataset is available in BIDS format via the Dutch research data repository DataverseNL at the following link: https://dataverse.nl/dataverse/rt-me-fmri. This repository includes the raw BIDS data, descriptive metadata, and derivative data including quality reports.

Apart from the dataset README file, all core files are available in one of 3 formats: NIfTI, TSV and JSON. Functional and anatomical data are stored as uncompressed NIfTI files (with the “.nii” extension), which contain image and header data and can be handled/viewed by all major neuroimaging analysis packages and programming languages. Tabular data such as participants, task events, response timing and physiology data are stored in tab-separated value text files (with the extension “.tsv”, or if compressed “.tsv.gz”) and can be handled by text or spreadsheet reading/editing software on all major operating systems, or alternatively by all major software programming languages. Metadata about the dataset, tasks, events and more are stored as key-value pairs in text-based JSON files (with the extension “.json”) that can be handled/viewed using all major software programming languages.

All data files are organised according to the BIDS convention for dataset participants, MRI data type (anatomical or functional), and derivatives, as depicted in Fig. 5.

**Figure 5:**
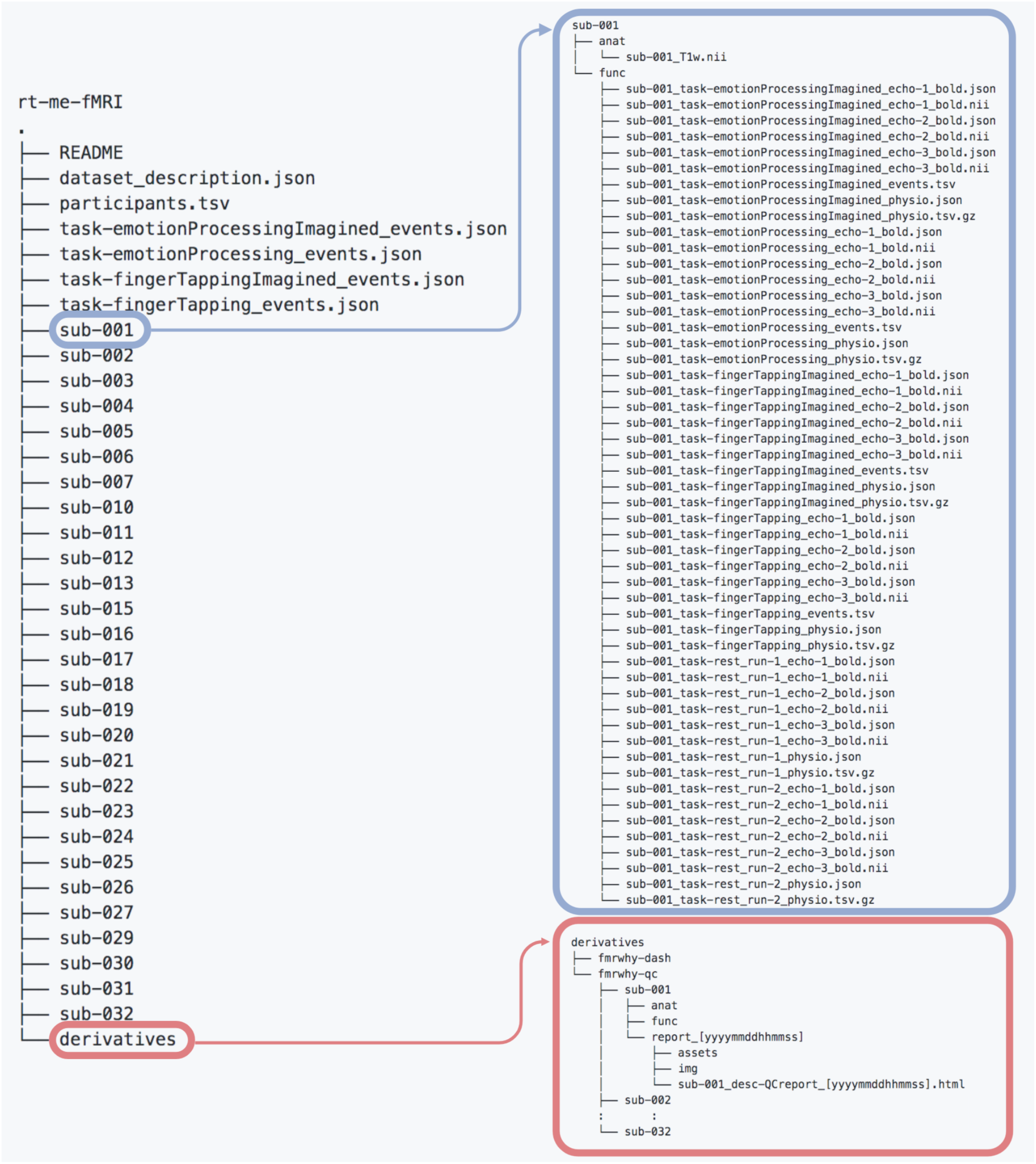
A diagram showing the content of the rt-me-fMRI dataset. The top level directory includes metadata about the dataset, participants and task events, as well as a directory per participant and lastly a derivatives directory. The expansion of “sub-001” (top right) shows subdirectories “anat” and “func”, each with neuroimages and metadata related to anatomical and functional scans, respectively. The expansion of the “derivatives” directory (bottom right) shows subdirectories “fmrwhy-dash” and “fmrwhy-qc”. The former contains all derivative data required to run the interactive browser-based application accompanying this dataset. The latter includes a quality report per participant in HTML format.

Each participant directory contains two subdirectories: “anat” and “func”, respectively containing all anatomical and functional images and metadata. Different data types can be distinguished based on BIDS identifiers, e.g. “_bold” for functional and “_T1w” for anatomical MRI data. The full list of data acquisitions with their data types, descriptions, and formats are provided below in Table 1. Note that for functional data, each resting state and task run consists of three separate image files, one per echo (i.e. “_echo-1_bold.nii”, “_echo-2_bold.nii”, and “_echo-3_bold.nii”). JSON sidecar files accompany all BOLD and physiology data files on the participant level, while the accompanying JSON sidecar files for the four types of task event files are on the dataset level. Other files on the dataset level include the README, the dataset description (JSON) and the participant list (TSV).

**Table 1:** *rt-me-fMRI* core dataset acquisitions, types, descriptions and formats.

| Acquisition | BIDS identifier (extension) | Data Type | Description | Acquired Data Format(s) | BIDS Format |
| --- | --- | --- | --- | --- | --- |
| T1-weighted | _T1w (.nii) | Anatomical MRI | Standard high-resolution | NIfTI | NIfTI |
| Resting data: task-rest_run-1 | _bold (.nii) | Functional MRI | Resting state | PAR/REC, DICOM | NIfTI |
| Task data: task-fingerTapping | _bold (.nii) | Functional MRI | Right-hand finger tapping | PAR/REC, DICOM | NIfTI |
| Task data: emotionProcessing | _bold (.nii) | Functional MRI | Matching shape and faces | PAR/REC, DICOM | NIfTI |
| Resting data: rest_run-2 | _bold (.nii) | Functional MRI | Resting state | PAR/REC, DICOM | NIfTI |
| Task data: fingerTappingImagined | _bold (.nii) | Functional MRI | Mental motor task - imagined finger tapping | PAR/REC, DICOM | NIfTI |
| Task data: emotionProcessingImagined | _bold (.nii) | Functional MRI | Mental emotion task - emotional memory recollection | PAR/REC, DICOM | NIfTI |
| Task responses and timing | _events (.tsv.gz) | Peripheral measure | Stimulus and response timing for all tasks, i.e. x4 | Eprime 'dat' and 'txt' files | TSV |
| Physiology data | _physio (.tsv.gz) | Peripheral measure | Cardiac and respiratory traces for all runs, i.e. x6 | Philips scanphyslog.log | TSV |
| n/a | n/a | Metadata | JSON sidecar files for all files of type _bold and _physio | Philips scanphyslog.log | JSON |

Apart from the core dataset, ***rt-me-fMRI*** also includes derivative data in two subdirectories generated by the *fMRwhy* toolbox and related scripts: “fmrwhy-qc” and “fmrwhy-dash”. The former results from the *fmrwhy_bids_workflowQC* pipeline and contains a subdirectory per participant, each in turn including subdirectories “anat” and “func”. These directories contain NIfTI, TSV and PNG files of quality control outputs, which are all required for the HTML quality report contained in the “report_[yyyymmddhhmmss]” directory. These reports can be opened with all major Internet browsers. The “fmrwhy-dash” derivative directory contains (as TSV files) all data required to yield the interactive visualisations of the supplementary browser-based application provided with this dataset: https://rt-me-fmri.herokuapp.com/.

## 4. Technical validation

### 4.1. BIDS validation

The full dataset was validated for BIDS compatibility with the use of the web-based “BIDS validator” tool (v1.5.4; available at https://bids-standard.github.io/bids-validator/). A log of the BIDS validator output can be found in the project’s code repository: https://github.com/jsheunis/rt-me-fMRI.

### 4.2. COBIDAS reporting

Data acquisition and experimental protocol parameters for this study were reported according to the community-formulated COBIDAS guidelines (Nichols et al., 2017). A modular version of this information is available in the project’s GitHub repository.

### 4.3. Data quality assessment

Image and data quality of this dataset was assessed using the *fMRwhy* toolbox. This allowed quality to be assessed for raw and minimally (pre)processed versions of the data, and also for interim steps on which the validity of eventual study outcomes might depend. A BIDS-compatible workflow in the *fMRwhy* toolbox, *fmrwhy_bids_workflowQC*, runs initial preprocessing and quality control of the raw data and outputs a quality report per subject, which includes metrics and visualizations for anatomical and functional MRI data and for peripheral data.

*For anatomical MRI:*

- Coregistered T1w segmentations (gray matter, white matter, CSF, and a whole brain mask) were overlaid onto the subject functional space, for visual inspection of the registration and segmentation quality.
- Coregistered anatomical regions of interest (in this case the left motor cortex, bilateral amygdalae and bilateral fusiform gyri) were overlaid onto the subject functional space, for visual inspection.

*For functional MRI (all runs):*

- A summary table provides values for all runs per subject for mean framewise displacement (FD), total FD, FD outliers, and mean tSNR in all tissue compartments. This allows quick inspection per participant, but is better understood when referenced to the whole dataset.
- Several image montages were generated per run, including the time series mean, the standard deviation and the tSNR map. The time series mean gives a quick view of the general quality of the time series and can indicate spike or interference artefacts. The standard deviation map shows areas with high signal fluctuation that can often be related to movement (e.g. close to the eyes). The tSNR maps are useful for investigating general signal quality, to indicate signal dropout and comparing signal quality across regions.
- A carpet (time series) plot was generated per run, which displays voxel intensity in percentage signal change from the mean over time. The vertical axis (voxels) is either grouped per tissue type (compartment ordered) or ordered from top to bottom according to the voxel’s time series correlation strength to the global signal. Signal traces above the carpet plot are also shown, including tissue compartment signals, respiration, heart rate, and framewise displacement. These plots are useful quality checking tools as they make it easy to visualise wide scale signal fluctuations across voxels, which can then be related visually to changes in physiological signals or subject movement.
- Checking the quality of the recorded cardiac and respiratory traces is made possible with images generated by TAPAS PhysIO during the process of calculating RETROICOR, CR and RVT regressors. Images include a plot of the temporal lag between derived heart beats within thresholds for outliers, and a plot showing the breathing belt amplitude distribution that can be inspected for unexpected shapes.

All functional quality metrics of the full dataset, generated by the *fmrwhy_bids_workflowQC* workflow, are summarised in Table 2. This includes, per run, mean framewise displacement, total framewise displacement, framewise displacement outliers (based on a conservative 0.2 mm threshold, and a liberal 0.5 mm threshold), and mean tSNR in all tissue compartments (grey matter, white matter, cerebrospinal fluid, whole brain). This allows possible data users to inspect the quality measures and to set personalised thresholds and exclusion criteria.

**Table 2: Functional quality metrics for the rt-me-fMRI dataset**

(Online version: https://github.com/jsheunis/rt-me-fMRI/blob/master/data/sub-all_task-all_desc-allQCmetrics.tsv)

Fig. 6 below displays summarised quality metrics for the rt-me-fMRI dataset, and examples of single-subject quality images. Individual quality reports can be downloaded together with the dataset.

**Figure 6:**
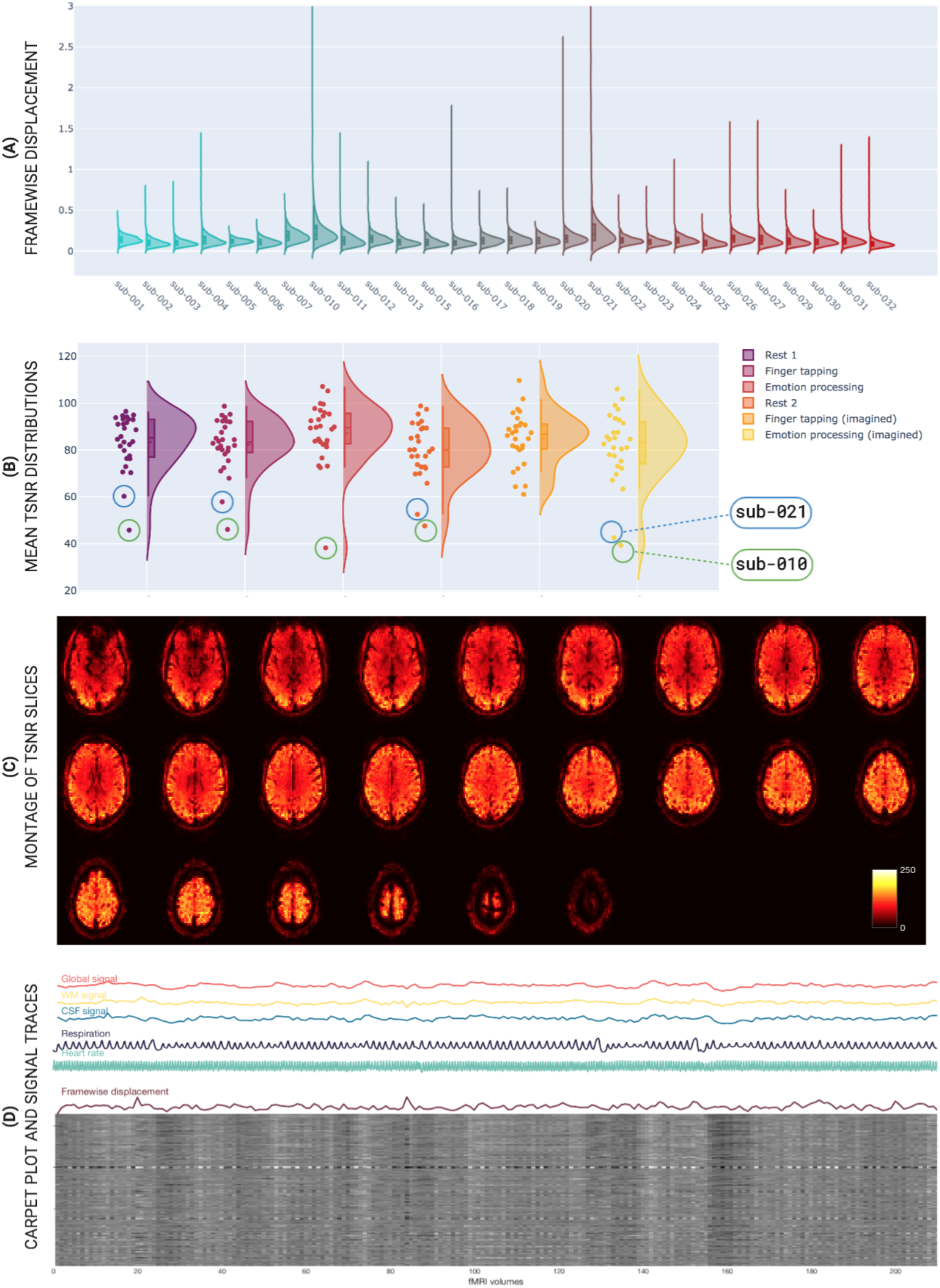
Representative quality checking information for the rt-mf-fMRI dataset. Subfigures include group level summary plots (A and B) and examples of subject level quality metric figures (C and D): (A) Vertical distribution (violin) plots of framewise displacement per subject, covering all functional runs. sub-010, sub-020 and sub-021 show comparatively high means and more outliers. (B) Vertical distribution (violin) plots of mean grey matter temporal signal-to-noise ratio (tSNR) per functional run, covering all subjects. Head movement results in higher signal fluctuations and hence lower tSNR, which is exemplified in the circled high mover data points: sub-021 (blue) and sub-010 (green); (C) An axial slice montage of temporal signal signal-to-noise ratio. (D) A time series “carpet plot” showing the global, white matter, CSF, respiration, and cardiac signals, as well as the calculated framewise displacement time series; (B) an axial slice montage of temporal signal signal-to-noise ratio.

### 4.4. Task validation

The slice timing corrected, 3D realigned and spatially smoothed Echo 2 time series of all task runs underwent individual- and group-level statistical analysis using a general linear model with SPM12. Task regressors included the main “ON” blocks for the *fingerTapping*, *fingerTappingImagined*, and *emotionProcessingImagined* tasks, and both the separate “SHAPES” and “FACES” trials for the *emotionProcessing* task. Regressors not-of-interest for all runs included six realignment parameter time series and their derivatives, the CSF compartment time series, and RETROICOR regressors (both cardiac and respiratory to the 2nd order, excluding interaction regressors, selected based on common implementation procedures in literature). Additional steps executed by SPM12 before beta parameter estimation include high-pass filtering using a cosine basis set and AR(1) autoregressive filtering of the data and GLM design matrix.

Contrasts were then applied to the single task-related beta maps for the *fingerTapping*, *fingerTappingImagined*, and *emotionProcessingImagined* tasks, and to the FACES, SHAPES, and FACES>SHAPES beta maps for the *emotionProcessing* task. Statistical thresholding, consisting of familywise error rate control with p < 0.05 and a voxel extent threshold of 0, was then applied on a per-subject basis to identify task-related clusters of activity. Unthresholded subject-level contrast maps were normalized to MNI152 space and then fed into a group-level one-sided t-test, for which the t-statistic maps were subsequently thresholded at p < 0.001 and an extent threshold of 20 voxels. Unthresholded individual- and group-level t-statistic maps can be accessed as a NeuroVault collection: https://neurovault.org/collections/XWDGUJHD/.

Fig. 7 below shows the resulting thresholded group t-statistic maps for all four task runs. Fig. 7A clearly shows activity clusters in the left motor cortex and right cerebellum, as expected for a finger tapping task as well as a negative activation pattern in the default mode network. Fig. 7C shows activation in the visual cortex commensurate with a face/shape matching task, specifically in the left and right fusiform gyri. Additional clusters are found in the amygdalae and hippocampi, as expected for an emotion processing task. For both imagined tasks, similar but weaker activation clusters are found in the expected regions (respectively the motor cortex in Fig. 7B, and amygdalae in Fig. 7D) but both wide scale activation patterns are consistent with mental tasks including imagery and memory recollection. Additionally, Figs. 7B and 7D show negative activation patterns in the dorsal attention network. The activation results in Fig. 7 are further evidenced by the resulting highest correlated terms when decoding the unthresholded group t-statistic images with the web-based Neurosynth tool (www.neurosynth.org, Yarkoni et al., 2011). Table 3 shows the resulting terms^2^.

**Table 3:** Neurosynth-decoded terms.

| Task | 10 highest correlated decoded terms |
| --- | --- |
| *fingerTapping* | *motor, premotor, finger, premotor cortex, movements, movement, hand, supplementary, execution, finger movements* |
| *emotionProcessing* | *face, fusiform, faces, fusiform face, fusiform gyrus, face ffa, ffa, occipital, inferior occipital, visual* |
| *fingerTappingImagined* | *theory mind, medial prefrontal, social, mind, mind tom, mental states, tom, primary, primary motor, junction* |
| *emotionProcessingImagined* | *medial, medial prefrontal, autobiographical, social, default, posterior cingulate, theory mind, mind, default mode, autobiographical memory* |

**Figure 7:**
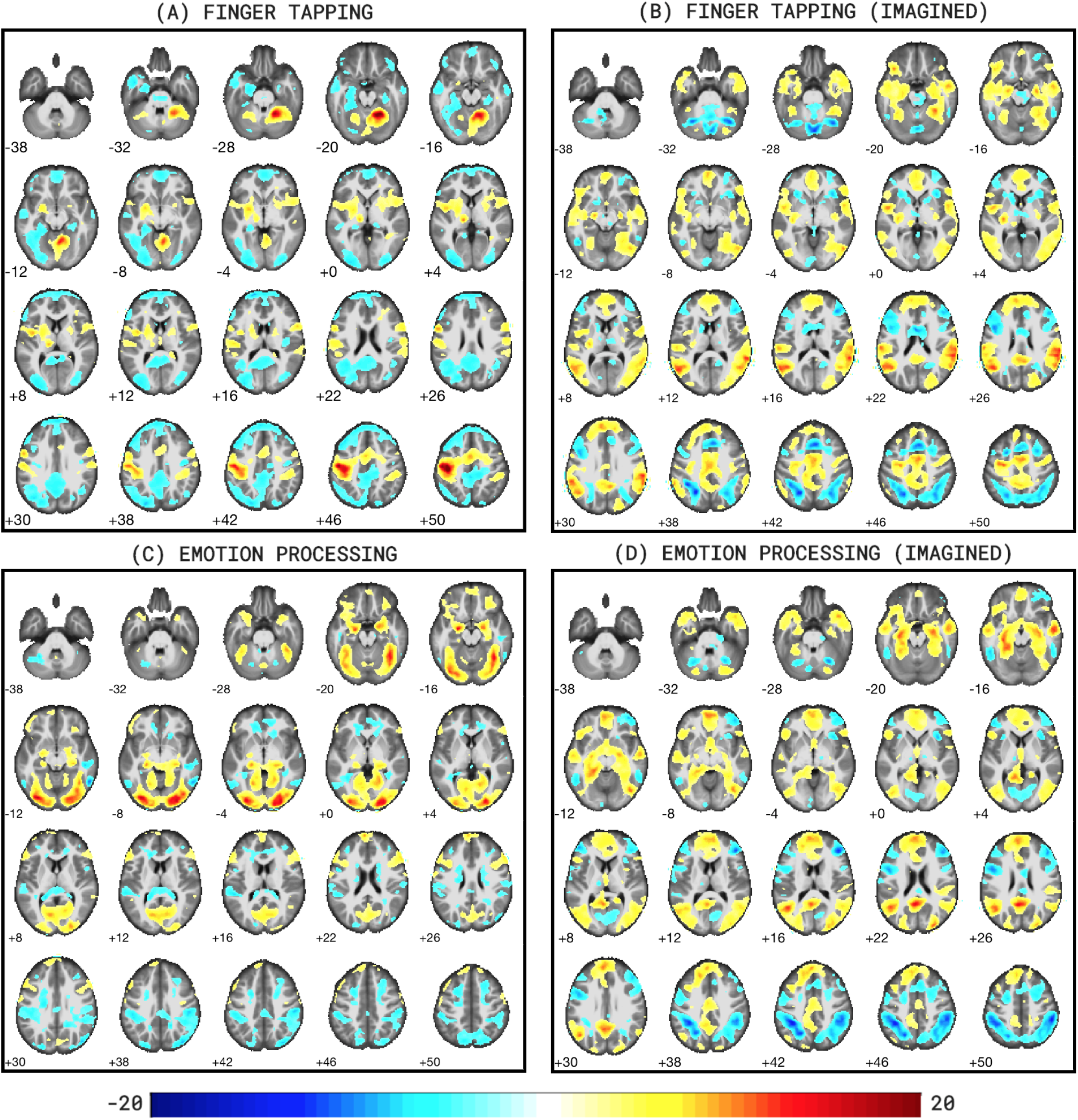
Group-level t-statistic maps for all tasks of the rt-me-fMRI dataset. Subfigures include: (A) fingerTapping, (B) fingerTappingImagined, (C) emotionProcessing, and (D) emotionProcessingImagined (p<0.001, voxel extent=20). Images were generated with bspmview. Fig. 7A clearly shows activity clusters in the left motor cortex and right cerebellum, as expected for a finger tapping task. Fig. 7C shows activation in the visual cortex commensurate with a face/shape matching task, specifically in the left and right fusiform gyri. For both imagined tasks, similar but weaker activation clusters are found in the expected regions (respectively the motor cortex in Fig. 7B, and amygdalae in Fig. 7D) but both wide scale activation patterns are consistent with mental tasks including imagery and memory recollection.

### 4.5. Multi-echo data validation

A core contribution of this ***rt-me-fMRI*** dataset lies in the multi-echo acquisition. Multi-echo fMRI samples multiple *T_2_*\*-weighted images at a range of echo times along the decay curve following a single transverse magnetic excitation, which theoretically allows the optimum BOLD contrast to be optimized for a range of baseline tissue *T_2_*\* values. Subsequently, echo combination through weighted summation or averaging is a typical processing step that generally increases temporal signal-to-noise ratio and contrast-to-noise ratio and decreases signal drop-out in regions with high susceptibility artefacts and signal dropouts (Menon et al., 1993; Posse et al., 1999; Posse et al., 2012). Echoes can be combined using a variety of weights, including baseline voxelwise tSNR and *T_2_*\* maps.

Figs. 8 and 9 illustrate that such combination procedures improve tSNR and signal dropout, hence validating the use of multi-echo fMRI for improved quality data. Representative signal recovery is demonstrated in the tSNR maps of Fig. 8 for a single run of a single subject, particularly by the blue and magenta arrows showing areas of signal dropout in the Echo 2 time series (including, respectively, the medial temporal and inferior temporal lobes, and the orbitofrontal lobe) and subsequent recovery in the combined time series. The light green arrows indicate substantial increases in tSNR in areas close to the bilateral temporal-occipital junction and towards the occipital lobe as the slices increase in a superior direction. Fig 9 shows distribution plots of the mean grey matter tSNR for the single (2^nd^) echo and two combined echo (tSNR-combined and *T_2_*\*-combined) time series, covering all functional runs and all subjects. The two combined echo time series clearly have improved tSNR values, increasing by ~30% from 85 (2^nd^ echo) to 112 (tSNR-combined).

**Figure 8:**
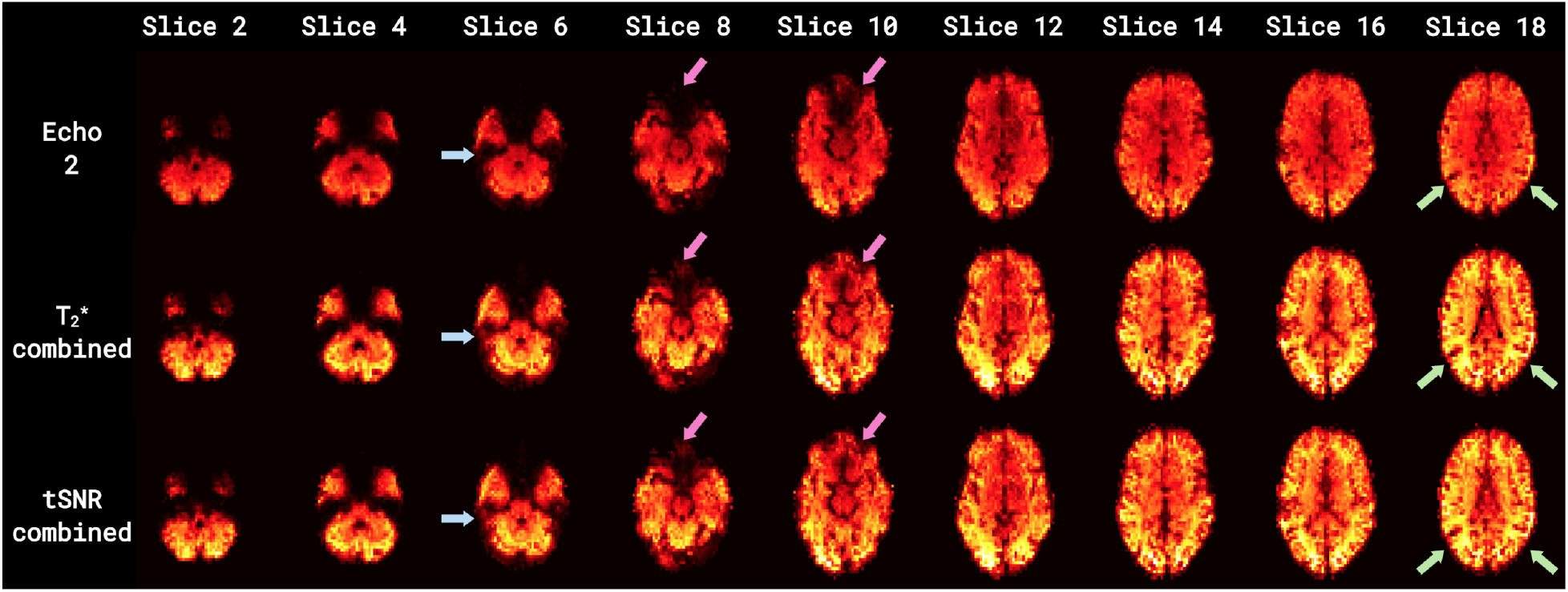
Axial slice montages of temporal signal-to-noise ratios (tSNR) in single and multi-echo combined time series. Time series include: the 2nd echo time series (top row) and two combined time series (middle row = T_2_*-combined; bottom row = tSNR-combined). Blue and magenta arrows indicate areas of signal dropout and recovery (including, respectively, the medial temporal and inferior temporal lobes, and the inferior frontal lobe). Light green arrows indicate areas with substantial increases in tSNR (in the lateral cortex and towards the anterior cortex as the slices increase in a superior direction). Combined multi-echo time series result in both substantially higher tSNR and signal recovery compared to Echo 2.

**Figure 9:**
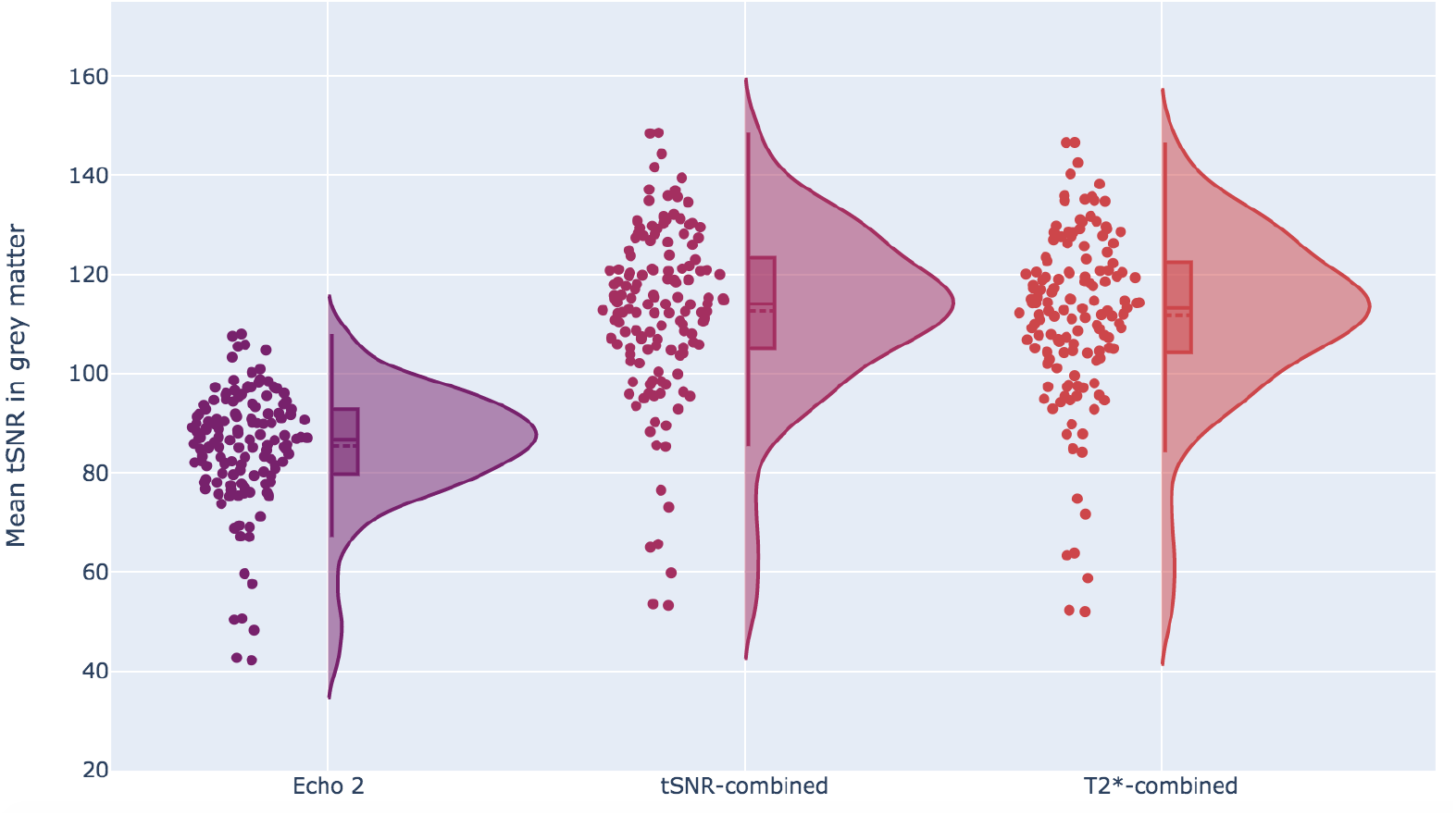
Vertical distribution (violin) plots of mean grey matter temporal signal-to-noise ratios (tSNR) in single and multi-echo combined time series. Distributions are shown for three time series: Echo 2, tSNR-combined and T_2_*-combined. A single distribution plot covers all subjects and all runs excluding rest_run-1. The two combined echo time series show clear increases in mean tSNR values.

Further benefits of multi-echo over conventional single-echo fMRI exist (see for example Olafsson et al., 2015; Dipasquale et al., 2017; Lombardo et al., 2016; Gonzalez-Castillo et al., 2016; Moia et al., 2020; Caballero-Gaudes et al., 2019), but such analyses are beyond the scope of this validation step and can be explored further with this publicly available dataset. In complementary work using this dataset we evaluate the use of several combination and *T_2_*\*-mapping procedures for both offline and real-time BOLD sensitivity (Heunis et al., 2020b).

### 4.6. Data inclusion/exclusion

To be a possible participant in this study, individuals had to be healthy, right-handed volunteers with no prior or current (at the time of the study) indications of neurological or psychiatric conditions. They also had to report the absence of any other standard contraindications for MRI scanning. 32 participants were initially recruited for the study, and the datasets of three participants were excluded due technical and one due to administrative challenges. No further datasets were excluded, even in cases of more than average or severe motion (e.g. sub-010 and sub-021), since it was decided that such data could still be useful for future methods development or related insights. Table 2 (also available in the project’s GitHub repository (https://github.com/jsheunis/rt-me-fMRI) provides a list of all functional quality metrics for all participants and runs, which allows possible data users to inspect the quality measures and to set personalised thresholds and exclusion criteria.

## 5. Code Availability

An interactive environment (https://rt-me-fmri.herokuapp.com/) was created alongside this study to allow users to interactively explore summaries of the data derivatives and quality control aspects.

All software scripts and self-developed tools used to prepare, preprocess and quality check the data are openly available at the project’s code repository https://github.com/jsheunis/rt-me-fMRI. This includes instructions to download, extract, and understand the data; the data preparation script; the preprocessing script; the quality reporting script; and the script to reproduce the figures for this manuscript.

Dependent software and toolboxes/packages used for these preparation, preprocessing and quality reporting steps include:

- *Python* 3.7+
- *bidsify* (v0.3; https://github.com/NILAB-UvA/bidsify)
- *scanphyslog2bids* (v0.1; https://github.com/lukassnoek/scanphyslog2bids).
- *dcm2niix* (v1.0.20190410; https://github.com/rordenlab/dcm2niix/releases/tag/v1.0.20190410)
- pydeface (v2.0.0; https://github.com/poldracklab/pydeface/releases/tag/2.0.0; Gulban et al., 2019)
- *convert-eprime* (v0.0.1; https://github.com/tsalo/convert-eprime/releases/tag/0.0.1; Salo, 2020)
- *MATLAB R2016b or later (9.1.0.441655; The MathWorks Inc)*
- *fMRwhy (v.0.0.1, https://github.com/jsheunis/fMRwhy)*
- *SPM12* (r7771; https://github.com/spm/spm12/releases/tag/r7771)
- *Anatomy Toolbox* (v3.0; Eickhoff et al., 2005)
- *bids-matlab (v.0.0.1, https://github.com/jsheunis/bids-matlab/releases/tag/fv0.0.1)*
- *dicm2nii* (v0.2 from a forked repository; https://github.com/jsheunis/dicm2nii/releases/tag/v0.2)
- *TAPAS PhysIO* (v3.2.0; https://github.com/translationalneuromodeling/tapas/releases/tag/v3.2.0; Kasper et al., 2017)
- *Raincloud plots* (v1.1 https://github.com/RainCloudPlots/RainCloudPlots/releases/tag/v1.1; Allen et al., 2019)
- *bspmview* (v20180918; https://github.com/spunt/bspmview/tree/20161108; Spunt, 2016)

## 6. Acknowledgements

This work was funded by the foundation Health-Holland LSH-TKI (grant LSHM16053-SGF) and supported by Philips Research. LH was supported by the European Union’s Horizon 2020 research and innovation program under the Grant Agreement No 794395. CCG was supported by the Spanish Ministry of Economy and Competitiveness (Ramon y Cajal Fellowship, RYC-2017-21845), the Basque Government (PIBA_2019_104) and the Spanish Ministry of Science, Innovation and Universities (MICINN; PID2019-105520GB-100).

## 7. Author contributions

SH designed the protocol, collected and curated the dataset, performed visual quality control, preprocessed and analysed the data, and wrote the article.

MB reviewed the article.

CC designed the protocol, contributed to data analysis, and reviewed the article.

LH designed the protocol, contributed to data analysis, and reviewed the article.

WH designed the protocol, contributed to data analysis, and reviewed the article

JJ designed the protocol, contributed to data analysis, and reviewed the article

RL designed the protocol, contributed to data analysis, and reviewed the article

SZ administered the project and reviewed the article.

BA administered the project and reviewed the article.

## Footnotes

1 Note: for the majority of participants, the presentation timing for the emotionProcessing task was delayed by tens of milliseconds for each trial (planned versus actual timing). This resulted in the full task presentation running on for about 5 s after the scan acquisition stopped. This is not deemed a problem, mainly since the exact presentation time was captured and is available in the BIDS dataset. However, users should take note not to use the planned timing parameters as that would ignore the delay that occured.

2 Task names of the **rt-me-fMRI** dataset were selected based on the desired activation response for the given use cases, e.g. emotionProcessing to elicit a response in regions involving emotion processing, with the knowledge that the tasks might yield varied responses and have varied use cases. This can lead to the activation analysis and Neurosynth decoding process yielding patterns and terms that do not necessarily reflect the task name, e.g. activation of the fusiform face area and related terms (“face”, “fusiform”, “occipital”) for the emotionProcessing task.

